# Complex molecular mixtures under cycling gradients as basis for life’s origins

**DOI:** 10.1101/050740

**Authors:** Jan Spitzer, Bert Poolman

## Abstract

We consider life as a cyclic physicochemical process that makes heredity and Darwinian evolution observable through living cells. We elaborate four principles that constrain current speculations about life’s emergence to natural processes driven by diurnal physicochemical gradients, primarily of temperature, water activity and electromagnetic radiation. First, Earth’s prebiotic chemical evolution is historically continuous with Darwinian evolution; second, cycling energies of solar radiation are primary drivers of chemical evolution; third, environmental molecular complexity must be high at the origin of life; and fourth, non-covalent molecular forces determine molecular recognition and cellular organization. Under normal physiological conditions of high ionic strength and high macromolecular crowding, hydration interactions (hydrogen bonding), screened electrostatic forces and excluded volume repulsions act over a *commensurate* distance of about one nanometer. This intermolecular distance governs chemical coevolution of proto-biomacromolecular surfaces (nucleic acids, proteins and membranes) toward Darwinian thresholds and living states. The above physicochemical principles of life’s emergence are consistent with the second law of thermodynamics, and with the current facts of molecular microbiology and planetary sciences. New kinds of experimentation with crowded molecular mixtures under oscillating temperature gradients - a PCR-like mechanism of life’s origins - can further illuminate how living states come about.

**Graphical abstract:** Life’s emergence follows from chemical and Darwinian evolution, a high degree of molecular complexity and a high crowdedness, and non-covalent molecular forces that determine molecular recognition and cellular organization. The macromolecules divide the cytoplasm into dynamically crowded macromolecular regions and topologically complementary electrolyte pools. Small ions and ionic metabolites are transported vectorially between the electrolyte pools and through the (semi-conducting) electrolyte pathways of the crowded macromolecular regions.

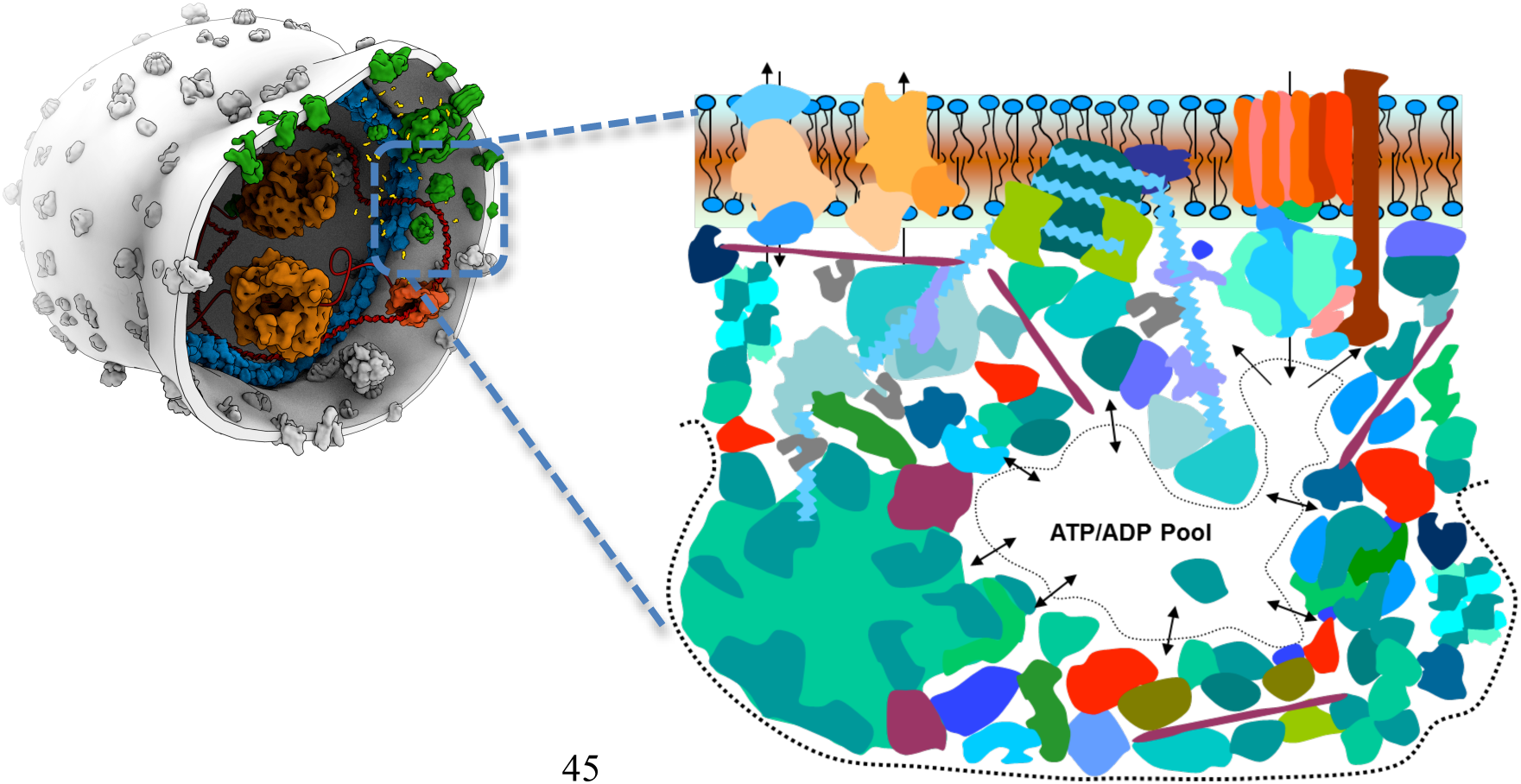

## Introduction

*By the ‘riddle of life’ not everybody will understand the same thing. We all, however, desire to know how life originates [in the universe] and what death is…* Jacques Loeb, 1912

The complexities of the origins and emergence of life remain unresolved. From the vantage point of cellular life, Franklin Harold reviews the state of research on life’s origins and finds it ‘in a limbo’. Over the last 60 years, he says, the research ‘has failed to generate a coherent and persuasive framework that gives meaning to the growing heap of data and speculation’ (Harold, 2014). Harold suggests that there is more to life than what physics and chemistry can explain - something that is eluding us and that may remain a mystery. Peter Atkins, from the standpoint of physical chemistry, also admits to the persisting enigma of life’s origins but sees it as a tough puzzle that will be solved by science, and by chemistry in particular - by establishing empirical facts and giving them deep theoretical meaning while rejecting models that do not work (Atkins, 2011). Atkins says that research on life emergence ‘is not stumped: it is alive with ideas but does not yet have sufficient [experimental] evidence to identify which of them, if any, is correct [at this time]’. Harold’s concern about the lack of a theoretical framework has been recently expressed also by a workshop on origin of life research that identified ‘an urgent need for a better, comprehensive theory of life to better define the aims of origin of life investigations…’(Scharf *et al.*, 2015); we add that any such general theory must be consistent with current physicochemical laws and with the facts of bacteriology and planetary sciences, and, importantly, it ought to suggest new kinds of experimentation, as outlined in this review.

Though Harold’s harsh critique of origins research seems justified, it is questionable that new research paradigms lie outside chemistry - in elusive molecular organization of cells that physics and chemistry cannot explain. In fact, from a physicochemical standpoint, there has been no shortage of ‘beyond chemistry’ concepts that attempt to explain molecular origins of living states. They involve unexplained appearances of self-replicating ribozymes that refuse to diffuse and mix with other prebiotic chemicals, of inexplicable self-assembly of abstract autocatalytic metabolic cycles, of indeterminate ‘flows of energy’ creating biological order, of assemblies of molecules endowed with Darwinian evolution or with the ability to arise from themselves, or endowed with autonomy and agenda bordering on free will and anthropocentric purpose; algorithmic ‘replicators’ and alien life based on non-carbon chemistries were also suggested, and many other ideas, assembled and reviewed in thematic collections and books (Deamer & Fleischaker, 1994; Lahav, 1999; Fry, 1999; Zaikowski *et al.*, 2008; Bedau & Cleland, 2010; Szostak & Deamer, 2011; Lane, 2015). As Harold intimates, the effort has been prodigious but understanding is lacking.

Peter Atkins brings to origins research the ‘old’ perspective of physical chemistry, partly reviving Jacques Loeb’s mechanistic conception of life (Loeb, 1962), by drawing attention away from the current preoccupation with DNA and RNA replication - from biological information. Tongue-in-cheek, Atkins says, ‘at a molecular [DNA] level, everything is junk…and we just happen to be [evolutionarily] a very successful junk’ (Atkins, 2011). Atkins’s observations on cellular complexity and genetic information have a deep physicochemical and evolutionary meaning for origin of life research. It implies that complex mixtures of molecules had to chemically evolve and structurally organize in order to cross the (phylogenetic) Darwinian threshold into hereditary cellular life that does carry genetic information (Woese, 1998; 2000; 2002). Here, Harold and Atkins seem to agree: no molecules carry genetic information - only living cells do so - and only with the help of many kinds of other cellular and environmental (nutrient) molecules.

Atkins’s physicochemical view brings into focus two (related) aspects of life’s emergence that have been neglected. First, there is the issue of thermal disordering effects of the second law of thermodynamics, manifested by ever-present diffusion (Brownian motion) without which life is impossible but which also drives cellular organization towards disorder, and eventually to physicochemical equilibrium (death). And second, the role of attractive and repulsive non-covalent forces that counteract molecular diffusion and guide the assembly of biomacromolecular complexes, thus enabling cellular life and delaying death. We whole-heartedly embrace Atkins’s emphasis on life being complex non-equilibrium chemistry that must follow thermodynamic laws and Harold’s view that life’s emergence can be considered only as ‘molecules into cells’ (Harold, 2005) - the spatiotemporal molecular organization of ‘first’ bacterial-like organisms, the populations of which gave rise to single-cell and multicellular eukaryotic organisms. We organize our physicochemical review on life’s emergence as follows. First, we put forward a working (biological) definition of life, and then describe generic physicochemical properties of cellular complexity derived from factual properties of bacterial cells. Second, we introduce and then discuss four physicochemical principles that constrain ideas about life’s emergence to cyclic physicochemical processes acting on multiphase and multicomponent chemical mixtures. We stress the importance of non-covalent intermolecular forces - and of electrostatic interactions in particular, which are fundamentally responsible for molecular recognition, the assembly of biomacromolecular complexes, and ultimately for the overall cellular organization. Based on such physicochemical principles, we suggest new kinds of experiments with cycling physicochemical gradients acting on complex molecular mixtures, which represents a proto-PCR mechanism of life’s emergence.

### Life as a cyclic (evolutionary) physicochemical process

Defining life has generated a large number of research communications that have a questionable usefulness (Szostak, 2012; Trifonov, 2012), though some clarifications are admittedly necessary (Benner, 2010; Cleland & Chyba, 2002; Luisi, 1998). We are skeptical about definitions of life that endow ‘dead’ molecules with non-chemical (biological) properties, such as the NASA definition of life as ‘a self-sustaining chemical system capable of Darwinian evolution’ (Joyce, 1994). This definition simply assigns Darwinian (biological) evolution to a mixture of molecules containing nucleic acids, something inadmissible from a physicochemical standpoint: no molecules can exhibit Darwinian evolution and besides, no self-sustaining molecular systems (e.g., simpler ones that are not capable of Darwinian evolution?), have been discovered, nor are they likely to be discovered. Thermodynamic laws mandate that non-equilibrium chemical systems left to themselves drift to physicochemical equilibrium (death), minimizing free energy in accordance with the first and second law of thermodynamics (Atkins, 2011). Thus, for example, the simplest manifestation of life related to issues of life’s emergence - a population of bacterial cells - dies off after the stationary phase of growth, reaching a physicochemical equilibrium; however, we note that it is no simple matter to verify the death of a bacterial cell (Davey, 2011; Siegele & Kolter, 1992).

The NASA definition of life is thus an unnecessary tautology that assigns biological concepts to lifeless molecules, indirectly endorsing the view that life’s molecules and macromolecules - DNA and RNA in particular, are in some sense special. According to this view, first promulgated by Schrödinger (Schrödinger, 1944), the precision of cellular reproduction (heredity) is so astonishing that, at the molecular level, it must involve new biophysical laws equivalent in scope to thermodynamics or quantum mechanics which endow biomacromolecules with special ‘biological’ or replicative properties. Since the appearance of Schrödinger’s book, however, cellular heredity became understood via biochemistry (structure and properties of the DNA double helix, genetic code, the ‘dogma’ of molecular biology, PCR protocols, genetic engineering, etc.), making it clear that at the molecular level DNA and RNA are ‘dead’ and incapable of generating living states (Atkins, 2011); *cf*. also Perutz’s review of Schrödinger’s book (Perutz, 1991). From a chemistry standpoint, we know that all molecules including biomolecules and biomacromolecules follow the same physicochemical laws and that their properties are independent of the methods of their syntheses - whether enzymatic (biological) *in vivo* or *in vitro*, or via unrelated synthetic steps of organic chemistry. Therefore, the ability of biomacromolecules to maintain a reproductive cellular organization lies in their non-equilibrium (cell cycle) chemistry in a given physicochemical (nutrient) environment, governed by existing laws of thermodynamics and quantum chemistry. It is noteworthy that after 150 years, Pasteur’s ‘microbiological law’ - all life only from life - remains valid also in molecular thermodynamic sense: bacterial life may not emerge (self-assemble) *spontaneously* from its molecules (Spitzer, 2014).

In order to proceed toward meaningful experimentation related to the emergence of life, we suggest a physicochemical definition of ‘first’ life as - a cyclic physicochemical process that makes heredity observable in ancestral (bacteria-like) cells that grow and divide in a sustaining mixture of chemicals. Equivalently, in biological language, the ‘first’ life is - a repeated cell cycle during which one bacteria-like cell yields two (very) similar but non-identical cells in a nutrient environment. Two corollaries of this definition help explain Darwinian evolution at a single cell organismal level. First, the non-identity of progeny (partial or incomplete heredity) guarantees Darwinian evolution through subsequent cell cycles by creating Darwinian variations on which natural selection can act. Second, a repeated perfect cell cycle that yields two (‘mathematically’) identical cells that are both identical to the original cell lacks Darwinian variations and cannot evolve. Nevertheless, the perfect cell cycle represents an imaginable (ideal) bacterial life that some bacterial species may approach in their behavior (living fossils in a chemostat?), just as some gases approach ideal gas law under some conditions.

The evolving populations of such ‘first’ bacteria-like organisms are referred to as LUCA - the last universal common ancestor (Woese, 1998), from which contemporary Bacteria, Archea and Eukaryota (single-cell and multicellular organisms) evolved over the last 3.5 billion years. Such Darwinian evolution, represented by a tree of life with a pre-LUCA complex chemical root system (O’Malley & Koonin, 2011; Doolittle, 1999), has underlying physicochemical mechanisms that can distinguish three kinds of cellular evolution: (i) *micro-evolution* arising from the errors in the replication of DNA within any one cell cycle during the growth of a population, (ii) *meso-evolution* arising from interactions between ‘dead’ environmental DNA and RNA (including viruses) and other cells (bacterial transformation and transduction, which evolved into bacterial competence exhibited by some contemporary bacterial species, and into laboratory protocols of genetic engineering), and (iii) *macro-evolution* arising from direct cell-to-cell interactions (e.g. in biofilms) that gave rise to more complex eukaryotic cells and multicellularity. The latter two kinds of Darwinian evolutions are not related to DNA replication, as they involve fusions and re-organizations of cell envelopes with concurrent melting and recombination of nucleic acids, driven by cycling temperature, water activity and other gradients - by natural processes akin to the protocols of genetic engineering and polymerase chain reactions. The contemporary bacterial cell cycle is thus an experimentally accessible end-point of the evolution of LUCA, Fig. 1, and we can reasonably assume that ancestral bacteria-like cells and contemporary bacterial cells share the same *physicochemical* attributes of complexity (defined in the next section). Therefore physicochemical conditions under which bacterial cells function today resemble those under which first bacteria-like cells emerged ~3.5 billion years ago (Spitzer & Poolman, 2009, Spitzer 2011; 2013; 2014; Spitzer *et al.*, 2015).

**Fig. 1.**
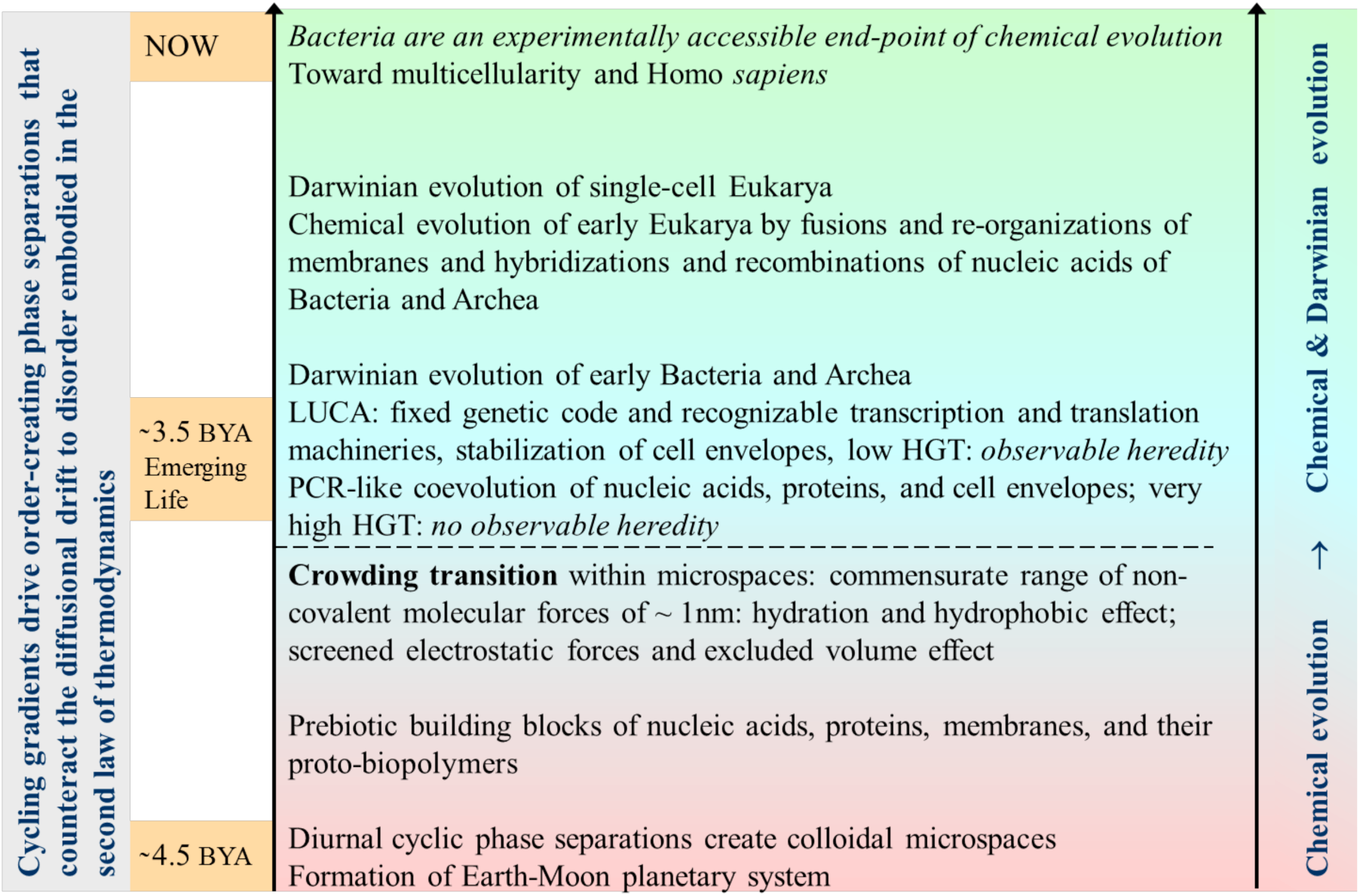
The historical continuity of chemical and Darwinian evolutions. The first living states probably appeared ~3.5 billion years ago (BYA). Contemporary bacteria are assumed to have similar complexity (biochemistry, genetics, and cellular structure) as the first ancestral bacteria.

### Cellular complexity

We take a broad view of Atkins’s lighthearted designation of nucleic acids as ‘molecular junk’, which cells utilize to exhibit short-term (generational, ‘Mendelian’) heredity and long-term (evolutionary, Darwinian) relatedness (Koonin, 2009), and re-define molecular ‘junk’ as complex chemical mixtures that comprise, in addition to nucleic acids, all other cellular biomolecules and biomacromolecules that make up a cell.

The chemical complexity of such mixtures has been rendered less mystifying over the last 200 years as cellular components were separated, purified, and their molecular structures, interactions and function determined - an astonishing success of the biochemical reductionist approach which has now culminated in molecular understanding of heredity via enzymatic replication of DNA double helix (Kornberg, 2000). In the current post-genomic era (Gierasch & Gershenson, 2009; Kell & Oliver, 2003; Eisenberg *et al.*, 2000), there is now enough established knowledge about all molecular components and physiology of microbial cells (Schaechter *et al.*, 2006; Kim & Gadd, 2008) that the reverse process of cellular reassembly - of ‘putting Humpty-Dumpty together again’, becomes conceivable. However, to re-construct the spatiotemporal molecular complexity of living cells - to re-create a living system from ‘dead’ biomacromolecules and other biomolecules - is a formidable task (Harold, 2005). Here, in the shadows of the unknown, broken and barely recognizable Humpty-Dumpty points to the need to ‘beat’ the second law of thermodynamics, as discussed later. We define generic physicochemical attributes of spatiotemporal complexity of bacterial cells as follows:

[1] *Phase separated* from the surroundings, *i.e.* bounded by a surface (interfacial, membranous) layers.
[2] *Multicomponent*, containing many kinds of small molecules, macromolecules, polyelectrolytes, ionic salts, and water.
[3] *Crowded*, with a high total volume fraction of macromolecules, which creates a system of *vectorial* electrolytic nano-channels that guide the diffusion of ions and metabolites (Spitzer & Poolman, 2005, 2009).
[4] *In disequilibrium*, both in chemical and physical sense, *i.e.* catalytically reacting (growing), and with physical inflows of water, ions and nutrients from the environment and vice versa
[5] *In a re-emergent process*, a ‘chemical engineering’ process that is cyclic (and evolving) with internal self-regulation that limits cellular growth by fission into two similar cells.

Conceptually, there is thus nothing cryptic about cellular complexity; rather than being irrevocably mysterious, it is a puzzle of too many kinds of molecules interacting together in a semi-liquid (sol-gel), electro-viscoelastic state maintained by non-covalent intermolecular forces and sustained by biochemical reactions that consume environmental nutrients and energies. Ultimately, the puzzle will be solved by taking bacterial cells apart and then putting the components back together, ensuring that the ‘assembly process’ is not thwarted by the diffusional drift to disorder - by the second law of thermodynamics (Spitzer, 2014).

The first four characteristics of chemical complexity define any *non-living* complex mixtures of molecules, e.g., water-based industrial formulations of paints, adhesives or inks drying under molecularly crowded conditions, which contain emulsion polymers (latexes) and inorganic insoluble fillers, such as calcium carbonate, together with functional chemicals such as buffers, thickeners, dispersants, coalescents, anti-foaming agents, anti-oxidants, etc. Non-exhaustive examples of molecular complexity related to the problem of origin of life include: the chemical matter of rotating planets and their moons (Bernstein, 2006; Carrasco *et al.*, 2009; Chyba & Sagan, 1992; Raulin *et al.*, 2012; Stoker *et al.*,1990), the great variety of chemical compounds (particularly those of carbon) which have been identified in the cosmos (Dworkin *et al.*, 2001; Rhee *et al.*, 2007; Ehrenfreund & Cami, 2010; Pizzarello & Shock, 2010), the readily formed ‘tars’ in non-enzymatic organic syntheses of prebiotic biomolecules and biopolymers (Miller, 1953; Shapiro, 1986) and contemporary corpses of biological origin in the process of reaching physicochemical equilibrium, including those of bacteria (Atkins, 2011; Davey, 2011).

The fifth property of re-emergence (cell cycle) is a unique property of living mixtures of molecules represented by contemporary bacterial cells. Re-emergence defines the current bacteriological problem of ‘being alive’ vs. ‘being dead’ and anything in between, a biophysicochemical state that is strongly dependent on environmental conditions. Re-emergence implies that only cycling (oscillating) chemical evolutionary processes could have led to the historical (prebiotic) emergence of life - to the first reproducing cells with a replicating DNA. In other words, continuous non-steady ‘random’ chemical processes (chance) are extremely unlikely to evolve into repeatable metabolic and genetic processes of a bacterial cell cycle. Incidentally, the physicochemical understanding of ‘being alive, dormant, sick or dead’ is also relevant to the problem of ‘unculturable’ bacteria believed to exist in large numbers in the environment but not yet grown in the laboratory, and to medical issues involving pathogenic bacteria (Davey, 2011; Bauermeister *et al.*, 201; Oliver, 2010; Bogosian & Bourneuf, 2001; Stewart, 2012).

Only when non-equilibrium complex molecular mixtures are continuously phase-separating and chemically reacting under *cyclic* non-steady state conditions, *i.e. repeatedly stoked with energy*, only then their chemical evolution into living states becomes conceivable. Only then, the diffusional drift to disorder, governed by the second law of thermodynamics, can be temporarily reversed and molecular chaos defeated, when intermolecular non-covalent forces come into play under crowded molecular conditions. In general, these molecular forces explain the existence of lower entropy chemical phases when we go from mixed gases and vapors to liquids, solutions, and sols - and other ‘semi-liquid’ colloidal phases - and to solid gels, amorphous solids, and crystals. Classical physical chemistry cannot deal with bacterial molecular complexity of ‘too many components and phases’ in the traditional reductionist manner of chemical thermodynamics and kinetics. Hence an empirical term ‘crowding’ was introduced to recognize a *total* high concentration of many cellular biomacromolecules *in vivo*, some of which may exist at low individual concentrations. Crowding has been demonstrated to modulate protein folding, protein-protein and protein-nucleic acid interactions *in vivo*, making the cell function ‘on the brink of phase separations’. In comparison, classical *in vitro* biochemistry deals typically with single purified biomacromolecules at low concentrations, away from phase transitions and unwanted interactions (Srere, 1985; McConkey, 1982; Ellis, 2001; Zimmerman & Minton, 1993; Wang *et al.*, 2011; Zhou *et al.*, 2008; Mika & Poolman, 2011; Sarkar *et al.*, 2013; Foffi *et al.*, 2013; Rowe, 2011; Record *et al.*, 1998; Monteith *et al.*, 2015; Boersma *et al.*, 2015). Thus further experimental progress in ‘putting Humpty-Dumpty back together again’ (Gierasch & Gershenson, 2009) - making life emerge from ‘dead’ molecules - is likely to be along empirical avenues with ‘crowded’ systems; they will be well-defined by new methods of preparation, e.g. from existing bacterial populations (Spitzer, 2014), and by new methods of analyses to characterize weak associations of crowded biomacromolecules, such as ultracentrifugation (Rowe, 2011, Schuster & Laue, 1994).

The importance of attractive and repulsive non-covalent molecular forces (hydrogen bonding, hydration and the related hydrophobic effect, screened electrostatic forces and excluded volume effect) for cellular organization represent one of our four tenets that constrain speculations about life’s emergence and evolution to a more rigorous physicochemical basis (Spitzer *et al.*, 2015); these tenets are summarized and discussed below.

### Toward a theory of life’s emergence

We formulate four principles that unite chemistry and biology at the origin of life, derived from the above descriptions of bacteria-like first cells and their physicochemical complexity. These principles constrain speculations about life’s emergence to evolving complex chemical systems driven by cyclic disequilibria. We emphasize their consistency with chemical thermodynamics, with the facts of planetary sciences, phylogenetics, molecular biology, and with the well-understood non-covalent intermolecular forces, which are ultimately responsible for cellular ‘self-construction’ (Harold, 2005).

#### 1. Earth’s prebiotic chemical evolution is historically continuous with Darwinian evolution.

The continuity between chemical and Darwinian evolutions represents the culmination of chemical evolution of complex prebiotic molecular mixtures into tangible bacterial-like cells, Fig. 1. From phylogenetics, we modify Woese’s concept of Darwinian thresholds (Woese, 1998; 2002; Koonin, 2009; 2011) to include the role of cellular envelopes. As the cellular envelopes became gradually more stable, horizontal gene transfer decreased sufficiently for cellular identity (heredity) to persist and thus become observable, which signifies the beginning of biology. Increased cellular stability came about by combination of proto-lipids, and proto-peripheral and proto-membrane proteins, together with attachments of proto-nucleic acids to the membrane. The latter enabled the evolution of heritable molecular transport and ion gradients across the cell envelope - a key requirement for the evolution of cellular homeostasis, including the management of osmotic disequilibria between the inside and the outside of a cell (Andersen, 2015; van den Bogaart *et al.*, 2007; Wood, 2015; Konopka *et al.*, 2009; Record *et al.*, 1998).

During the more ‘primitive’ (non-hereditary) stages of chemical evolution, the enzymatic replication of proto-nucleic acids was inefficient and their evolving meltings, hybridizations and re-combinations were strongly dependent on external cycling temperatures - a cycling process that is confirmed in current PCR protocols and by the physical chemistry of DNA helices, e.g., the dependence of DNA melting (unwinding, dissociation) and hybridization (re-winding, association) on temperature and ionic strength (Marmur & Doty, 1962; Schildkraut & Lifson, 1965). Similarly, in vitro reconstitutions of ribosomes require specific temperature manipulations and buffered ionic strength (Traub & Nomura, 1968; Sykes & Williamson, 2009), which can be regarded as a relic of their chemical evolution into contemporary nucleoprotein complexes (Hud, et al., 2013; Petrov, et al., 2015) driven by cycling temperatures in high ionic strength electrolyte.

#### 2. Cycling energies of solar radiation are primary drivers of chemical evolution.

Rotating Earth converts solar energy (Rothchild, 2003) into cycling physicochemical gradients that keep chemistry along Earth’s surfaces in cycling disequilibria; this cyclicity represents the fundamental process of prebiotic chemical evolution of early Earth. The diurnal gradients of electromagnetic radiation, temperature and water activity bring about order-creating colloidal phase separations (compositions of lower entropy compared to the more random environments), representing a physicochemical mechanism of formation of microspaces - the chemically evolving precursors of cellular envelopes of ancestral bacterial cells. Such cyclic phase-separations continuously counteract the diffusional drift of prebiotic molecules to disorder mandated by the second law of thermodynamics. Cyclic colloidal phase-separations and are thus an integral part of physicochemical processes that evolved into cyclic living systems.

#### 3. Molecular complexity must be high at the origin of life.

Only a subcategory of chemicals can cyclically phase-separate from Earth’s total physicochemical complexity, *i.e.* keep appearing and disappearing, and thus evolving with time as colloidal structures with permeable boundaries, while the environment outside is becoming a reservoir of nutrients. Only when Earth’s atomic composition is favorable for the formation of future biomolecules (under the prevailing conditions of cycling temperatures, radiation and water activities), the evolution of colloidal microspaces into cycling living states becomes conceivable. Thus the chances of chemical evolution toward living states are maximized when the atomic (ionic) and molecular diversity of early Earth is large. Earth’s molecular complexity, originating from diurnal disequilibria in Earth’s atmosphere, hydrosphere and lithosphere, is further expanded with the geochemistry of hydrothermal vents (Martin *et al.*, 2008) and with the astrochemistry of asteroids, meteors, comets and interplanetary dust particles that fall into Earth’s atmosphere (Chyba & Sagan, 1992; Bernstein, 2006; Rhee *et al.*, 2007; Pizarrelo, 2010). Taken together, these non-equilibrium processes result in a great variety of environmental chemical compounds comprising the elements of C, H, O, N, S, and P, dissolved and suspended in a complex electrolyte (seawater). These compounds never reach physicochemical equilibrium on account of the cyclic diurnal gradients of temperature, water activity and electromagnetic radiation, *i.e.* they keep on chemically evolving.

#### 4. Non-covalent molecular forces determine molecular recognition and cellular self-organization.

A neglected aspect of physicochemical complexity of extant bacterial cells (dead or alive) is the high volume fraction of all molecules within, which has been described as biomacromolecular ‘crowding’. It is now well established that proteins and nucleic acids are crowded within biological cells and permeated by a complex aqueous solution of dissolved small ions and molecules (metabolites), which allows for the evolution of metabolic and genetic pathways via molecular recognition and cellular organization. Fundamentally, spatiotemporal molecular recognition and cellular organization are determined by biochemical reactions and by non-covalent chemical interactions (Spitzer & Poolman, 2009; Spitzer, 2011; Parry *et al.*, 2014). Out of many non-covalent intermolecular forces (Dill, 1990), we find that a combination of four kinds - hydrogen bonding and hydration (Pauling & Corey, 1956; Eisenberg, 2003; Pal *et al.*, 2002; Park *et al.*, 2008), the related hydrophobic effect in aqueous media (Southall *et al.*, 2002; Rose & Wolfenden, 1993), screened electrostatic forces of the Debye Hückel type (Schildkraut & Lifson, 1965; Spitzer, 1984, 2003; Spitzer & Poolman, 2005), and excluded volume effect (crowding) - have a *commensurate* distance of action of about one nanometer, ensuring their *joint* participation in chemical evolution of biomacromolecular surfaces (Spitzer & Poolman, 2009; Laue, 2012). High biomacromolecular crowding in particular is a fundamental condition for life’s emergence, as it gives rise to transient vectorial channels within the gelled fraction of the cytoplasm - to complex vectorial biochemistry adjacent to the cytoplasmic side of the membrane, Fig. 2, and thus vectorially connected to membrane channels, transporters and other membrane proteins, *i.e.* to the environment (Spitzer & Poolman, 2005, 2009, 2013; Spitzer, 2011). The implications of these four principles for origins research are further elaborated and discussed below.

**Fig. 2.**
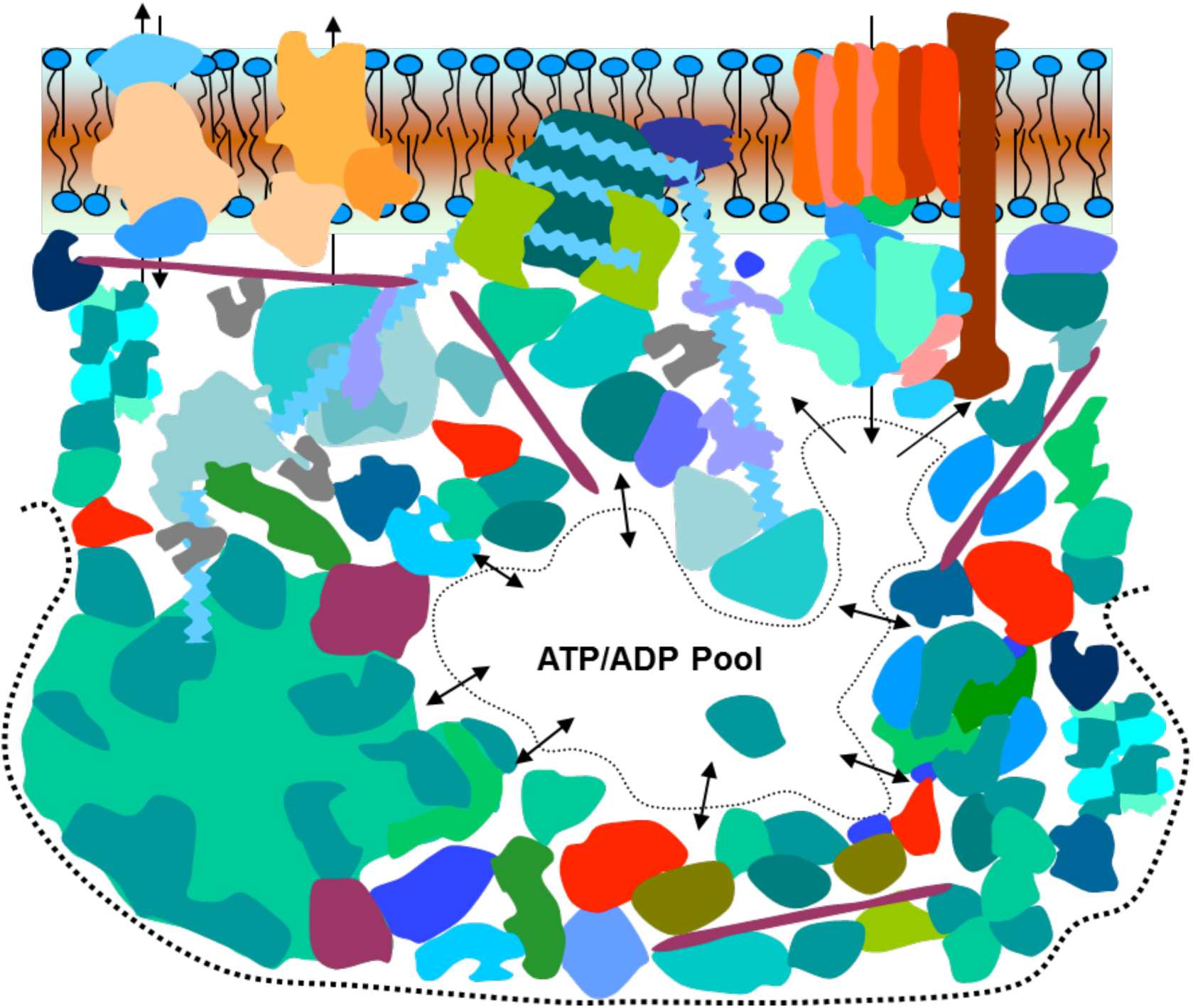
A cartoon of spatiotemporally supercrowded biomacromolecules and quaternary complexes, ‘spot-welded’ by attractive non-covalent forces, creating a gelled multiplex of electrolyte/metabolite pools and semi-conducting channels (complex vectorial biochemistry). The channels separate and localize ATP and ADP(AMP/P) and other ions according to their net ionic charge into different parts of cytoplasm, while making vectorial connections with membrane-located proteins, including ATP-synthase (top right-hand). Reprinted with permission from (Spitzer, 2011).

### Historical context and discussion

Traditionally, research on life’s origins has been based on Oparin’s and Haldane’s hypotheses reprinted in Bernal’s book (Bernal, 1967), which assume that living cells arose naturally on early Earth from prebiotic molecules and macromolecules (a ‘primordial soup’). Oparin’s and Haldane’s hypotheses are a direct response to Darwin’s theory of evolution, which left the question of the ‘first life’ unanswered. These hypotheses exclude the possibility of external agents of various degrees of omniscience and omnipotence to ‘seed life’ on Earth in one way or another. Astrochemical and planetary observations, as well as continuing ‘plausible’ prebiotic organic syntheses motivated by Stanley Miller’s electrical sparking of prebiotic atmospheres (Miller, 1953; Saladino *et al.*, 2012; Patel *et al.*, 2015; Dworkin *et al*, 2001), have now established that the Universe is capable of generating essentially all building blocks of life (low molecular weight amino acids, sugars, nucleobases, phosphates, and their organic derivatives) but *only* in complex high entropy molecular mixtures with many other environmental carbon compounds, including hard to characterize oligomeric and macromolecular compounds of tarry character with unsaturated and aromatic fused carbon rings, similar to tholins observed on Saturn’s moon Titan (McDonald et al., 1991; Carrasco et al., 2009). *How could life emerge from such prebiotic complex chemical mixtures in a natural way?* We have taken a physicochemical viewpoint (Spitzer & Poolman, 2009) summarized in our four principles (Spitzer *et al.*, 2015) that thermodynamically constrain the wide range of ideas about life’s emergence to cyclic processes of chemical phase-separations.

#### The continuity of Earth’s chemical and Darwinian evolutions

The chemical evolution of inanimate complex molecular mixtures and of living (Darwinian) states are separate but continuous - there is no discontinuous (‘miraculous’) transition (or dichotomy) between such states, e.g., ‘life being breathed’ somehow into molecular mixtures, or life being somehow ‘seeded’ on Earth by external agents. Neither can we assume that molecules synthesized enzymatically in living cells (particularly nucleic acids) are in some sense special, having evolved from primordial ‘replicator’ molecules in ‘simple protocells’ according to elusive biophysical ‘super-laws’ (Schrödinger, 1944; Pascal & Pross, 2015; England, 2013), which somehow banish diffusion (and thus entropic mixing), and so subsume the second law of thermodynamics in an apparent paradox. Given the established interpretations of chemical thermodynamics and quantum chemistry, the non-equilibrium association of biomacromolecules into living states lies within the current understanding of non-covalent intermolecular forces, the most important of which are hydrogen bonding (hydration) and the related hydrophobic effect, excluded volume repulsions, and screened electrostatic forces.

Our view of Darwinian thresholds is somewhat different from Woese’s, who assumed fixed (stable) genetic code - representing a threshold at which progenotes became ‘genotes’, with subsequently evolving transcription and translation molecular machineries. More likely, the threshold was a gradual transition defined by the increasing stability of cell envelopes (membranes) that minimized horizontal gene transfer and thus allowed cellular heredity to become observable, Fig. 1. Thus pre-prokaryotic chemical evolution of precursors of nucleic acids, proteins and cell envelopes were concurrent (chemically interacting), or at any rate could not get ‘too much’ out of phase in order to effectively evolve into cellular ‘first’ life (Spitzer, *et al.*, 2015). This physicochemical co-evolution model is indirectly supported by a recent synthetic scheme of organic prebiotic reactions driven by UV light, which can account for precursors of nucleic acids, proteins and lipids from a single carbon source of hydrogen cyanide (Patel *et al.*, 2015). However, prebiotic reaction schemes of organic chemistry cannot by themselves account for the *natural* emergence of life, because they neglect the diffusional mixing of prebiotic molecules - they do not ‘defeat’ the second law of thermodynamics in a natural way. Nevertheless, ‘prebiotic’ organic syntheses do establish the vast reactive potentialities of early Earth, which is cyclically kept out of equilibrium by diurnal gradients. For instance, the major product of Stanley Miller’s experimentation with prebiotic atmospheres were ‘tarry’ substances (Shapiro, 1986), which did not seem of interest, even though they might act as confining water-insoluble proto-membranes, filled and permeated by a variety of evolving aqueous primordial ‘soups’ - a generic example of evolving microspaces; such microspaces could arise even in interstellar (pre-cometary) ices (Dworkin *et al.*, 2001).

#### The necessity of cycling environmental energies

Cycling external energies are necessary for chemical evolution, *i.e.* to keep complex molecular mixtures out of equilibrium and hence evolving (chemically reacting and phase separating). A ‘random’, non-cyclic application of external energies (chance) is extremely unlikely to bring about living states; in fact, it would contradict the requirement of evolutionary continuity, endorsing a ‘miraculous’ appearance of living protocells from unremarkable molecular mixtures. Crucially, cycling external energies are required to overcome the second law of thermodynamics by cyclic phase separations of microspaces of lower entropy from high entropy mixtures of environmental chemicals. The multicomponent nature of such phase separations (containing molecules of variable water solubility and hydrophobicity) strongly suggests that new phases will appear on colloidal microscales of 10 - 10,000 nm. Such new phases have been also investigated from a constructive designed standpoint, either as two-phase systems (Keating, 2012) or coacervates (Tang, et al., 2014).

The response of microspaces to diurnal temperature cycles can be ‘instantaneous’ (in phase, maintaining thermal and water activity equilibria with the cycling environment), or delayed (out of phase) when some processes take longer (hours rather than seconds) to become equilibrated. This latter case is of particular importance for chemical evolution toward cellular life, as the system then begins to retain some ‘memory’ (structural, chemical) from the past cycles, when, for instance, the rates of dissolution are slower than the rates of precipitation. In other words, the system begins to maintain an evolving memory of its chemistry and structure. Large or unusual changes in environmental conditions (e.g., impact of asteroids) may entirely destroy any evolving microstructures (and create new ones of different kinds), but the cyclic evolution of prebiotic colloidal microspaces is inexorable: it continues as long as the Sun irradiates rotating Earth (and billions of suns irradiate billions of rotating exoplanets everywhere in the Universe).

#### The necessity of high molecular diversity at the emergence of life

Molecular complexity of Earth’s environments must be high for more structured (lower entropy) compositions to phase separate as microspaces with *still sufficient* chemical complexity to enable confined proto-biochemical evolution, Fig. 1. Thus the initial proto-biochemical evolution was directly dependent on cycling temperatures and other physicochemical gradients, which represents a proto-PCR mechanism of confined coevolution of genetics and metabolism, broadly consistent with the coevolution theory of the genetic code (Wong, 2005). The confining surfaces of microspaces can be of inorganic or carbon chemistries, e.g., phase-separated tholin-like partially hydrolyzed proto-biofilms rich in carbon and hydrogen without distinct cells, or ‘hatcheries of life’ in the form of inorganic membranous vesicles (Segré *et al.*, 2001; Dworkin *et al.*, 2001; Martin *et al.* 2008), or anything in between these two extremes. From an experimental standpoint, the dynamic supramacromolecular (colloidal) structures of the prebiotic microspaces are largely unknown (Spitzer & Poolman, 2009; Spitzer, 2013), because they evolve by cyclic fractional precipitations and dissolutions from multicomponent electrolyte solutions containing other molecules of varying molecular weights, hydrophobicities and solubilities.

Earth as a global chemical reactor (Spitzer & Poolman, 2009; Stüeken *et al.*, 2013) has three sources of complex chemical disequilibria: those driven cyclically by solar radiation impinging on a rotating lithosphere, hydrosphere and atmosphere, which are supplemented by geochemical reactions of superheated seawater with hot magma at hydrothermal vents and by random in-fall of astrochemicals. Physicochemical gradients at hydrothermal vents are unidirectional (hot to cold) and cannot plausibly evolve and convert themselves into evolving cyclic processes (Yellowstone’s ‘Old Faithful’ notwithstanding). Importantly, hydrothermal vents increase the chemical complexity of the ocean by providing metal ions, such as magnesium and calcium, including transition metals, e.g., iron, zinc or molybdenum. The relevant multivalent ions can become chelated (in Werner type of coordination complexes) with dissolved prebiotic molecules, derived from HCN and formamide (Saladino *et al.*, 2012; Patel *et al.*, 2015), or with polyphosphate anions if available; this is a well-established mechanism that keeps multivalent metal ions in solution or in colloidally stable particles (depending on concentration and pH), and thus molecularly available for cyclic prebiotic chemical evolution.

#### Non-covalent molecular forces regulate complex physiological processes

We find that relevant non-covalent intermolecular, hydrogen bonding, hydration and the related hydrophobic effect in aqueous media, screened electrostatic forces and the excluded volume effect, act over a commensurate range of distances of around one nanometer (Spitzer & Poolman, 2009). The commensuration principle is derived from the facts that hydration forces act over 2 - 3 water-molecule diameters, screened electrostatic forces act over a little less than one nanometer at physiological ionic strengths, and biomacromolecular crowding ~ 25% (observed in living cells) separates the surfaces of average proteins also by about one nanometer. Thus biochemical evolution can take place only in crowded systems at relatively high ionic strength, when hydration (and related hydrophobic effect), screened electrostatic forces and excluded volume effect act jointly over the distance of about one nanometer. This commensurate distance is only weakly dependent on temperature (Spitzer & Poolman, 2009), which is consistent with microbial evolution over the entire liquid range of water - from freezing to boiling. The desirability of crowdedness for the emergence and evolution of biomacromolecules has been recognized before (Zimmerman & Minton, 1993, Orgel, 2004).

The description of cellular complexity is made more complicated by the fact that during the cell cycle there are about 1000-2000 concurrent and sequential biochemical reactions within the cell and between the cell and the environment. These biochemical reactions are rather well synchronized and regulated to yield physiological processes, such as: (i) the sensing of the extracellular environment and the import of nutrients into the cell by the cell envelope, (ii) the conversion of the extracellular signals into cytoplasmic signals and their reception by the cytoplasmic side of the membrane and by the nucleoid (iii) biosynthesis of low molecular weight ‘building blocks’ including ‘fueling molecules’ such as ATP and GTP (iv) activation/deactivation of constitutive membrane proteins for immediate responses to environmental inputs, (iv) gene activation, silencing, and transcription, (iv) biosynthesis of ribosomes (v) translation via ribosomes including insertion of proteins into the membrane, (vi) the initiation, control and termination of the enzymatic replication of the nucleoid and plasmids, and (vii) the cell division and other morphological movements (shrinkage, invagination, budding, adhesive gliding, sporulation, etc.). All these processes take place within the membrane (cell envelope) and cytoplasm, and, from a physicochemical standpoint, their dynamic self-organization is ultimately determined by non-covalent intermolecular forces, among which electrostatic forces play a dominant role.

There can be little doubt that coulombic (electrostatic) interactions and electrolytic semi-conduction play a major role in regulating the multitude of inter-linked physiological processes on a global cellular scale; this is so because ‘naked’ electrostatic forces (Coulomb’s law) are both very strong (comparable to covalent bonds) and very long-range compared to all other non-covalent molecular forces, their strength decaying with the square of the distance. The cell *must* (and does) operate in an aqueous electrolyte of a relatively high ionic strength in order to shorten the range of naked coulombic forces (eq. 1), and thus make them commensurate with other non-covalent molecular forces (especially with hydration and excluded volume repulsions) on a scale of a little below one nanometer (Spitzer & Poolman, 2009). The Debye-Hückel theory thus modifies Coulomb’s law approximately by the exponential term *exp(-κr)* as

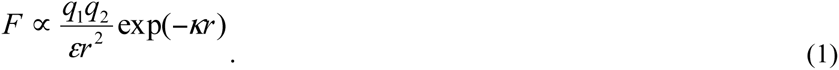

Here, *κ* is the Debye constant, the inverse of which, *1/κ*, is the Debye length, which is a measure of the effective range of screened electrostatic interactions: the higher the ionic strength, the faster screened electrostatic forces decay. At usual physiological ionic strength, the Debye length is a little below one nanometer. Thus, for a protein of radius 2.5 nm, the potential at 0.0001 molar salt is about 12 higher at its surface than at 0.1 molar salt; farther away at distance of 5.0 nm (2.5 nm from the surface) the potential is about 150 times higher, making the ‘self-assembly’ of biomacromolecular complexes at low ionic strengths much harder or impossible on account of stronger long-range electrostatic repulsion. The unique temperature dependence of the (high) dielectric constant of water makes the Debye length essentially independent of temperature, a fundamental circumstance that selects water as the biological solvent - there are no other solvents with similar dielectric constant so readily available in the Universe, providing an environment ‘fit for life’ (Henderson, 1913; Spitzer & Poolman, 2009; Spitzer, 2011). By the same token, the temperature independence of screened electrostatic interactions makes the emergence of life also almost independent of temperature, tremendously increasing the chances of chemical evolution toward living states.

It is not coincidental that when a bacterial cell is stressed (e.g., by starvation), the electrostatically highly charged molecules of (p)ppGpp are synthesized in order to bring very strong electrostatic forces into play within the cytoplasm, ‘ring the alarm bell’ so to speak, which initiates the *stringent* survival response on a global cellular scale. The response includes progressive shutting-down of transcription and of ribosome biosynthesis, the re-switching of genetic circuits, the densification of the nucleoid, the shrinking of the cell volume, etc. (Potrykus and Cashel, 2008; Siegele & Kolter, 1992; Frenkiel-Krispin *et al*., 2004). In order for such a complex chemical system to work reasonably reproducibly as it does during the cell cycle and respond successfully to environmental stresses, in other words, to effectively defeat the second law of thermodynamics through each cycle - biochemical reactions have to be localized and their localizations reproduced during each cell cycle; the localizations are reproduced by non-covalent intermolecular forces. Thus biomacromolecules are localized within the cell envelope, possibly as 2-D microdomains in the membrane (López & Kolter, 2010), including chemotaxis signal transduction complexes (Bray *et al*., 1998), and in ‘supercrowded’ 3-D micro-gels within the cytoplasm, as hypothesized by the sol-gel model (Spitzer & Poolman, 2013). In this model, ‘supercrowded’ (over 50% volume fraction of biomacromolecules) micro-gels are associated mainly with the cytoplasmic side of the cell envelope, and the nucleoid is situated in the middle with extensions into the cell envelope (Spitzer, 2011; Spitzer & Poolman, 2013). A similar qualitative interpretation was given to protein diffusion data in mitochondria, which describes cytoplasmic environment as crowded but ‘watery’, allowing for fast diffusion of small molecules (Partikian *et al*., 1998; Mika & Poolman, 2011). The cytoplasmic supercrowding extends the vectorial biochemistry of membrane proteins deeper into the cytoplasm by up to 70 nm, where the supercrowded microgels contain physical ‘microfluidic’ channels, the permeability and structure of which are controlled by biochemical reactions, Fig. 2. The channels have tunable structure controlled by electrostatic potentials, e.g., by phosphorylations (Spitzer & Poolman, 2005, 2013; Spitzer, 2011), which guide the important ‘energy’ ions ATP and GTP, and their less charged precursors (ADM and AMP, and other ions) according to their net charge through different parts of the gelled cytoplasm. This physicochemical mechanism (membrane and cytoplasmic ‘microfluidics’) represents complex vectorial chemistry of the cell, which limits and controls the diffusional disorder arising from the second law of thermodynamics - a key characteristic of living states.

The biomacromolecular gel formation (supercrowding) and the reverse process of gel liquefaction are controlled by biochemical signaling reactions that increase or decrease hydrophobic and screened electrostatic interactions, *i.e.* epigenetic modifications or ‘processing’ of nucleic acids and proteins during or after their biosynthesis. For example, methylations and de-phosphorylations increase non-covalent attractions via the hydrophobic effect and via reduced electrostatic repulsions, leading to gel formation; the reverse reactions, demethylations and phosphorylations, liquefy the gel. Intrinsically disordered proteins (IDPs), being more water-soluble and approaching the state of random coil on dilution, may also contribute to the dynamics of sol-gel transitions, particularly in respect to the ‘under-crowded’ regions of the cytoplasm (Bray, 2005; Theillet *et al*., 2014; Spitzer & Poolman, 2013; Spitzer, 2011). Cellular volume and viscosity changes associated with sol-gel transitions, along with the forces generated by localized membrane reactions that charge and discharge the membrane contribute to the morphological (bio-electromechanical) movements of the cell *in toto*; the membrane acts as a (‘leaky’) electrical viscoelastic capacitor, maintaining variable membrane potential and creating a multitude of chemiosmotic signals on the cytoplasmic side of the cell envelope (Mitchell, 1979). Some of these signals are transmitted ‘tangentially’ along the cell envelope to control (activate or deactivate) various transporters including the ATP-fed motion of the flagellum, and some are transmitted to the nucleoid to regulate gene expression, and thereby cell’s growth and division.

### New experimental outlook

The current experimental paradigm of origins research is based on the assumption of complexification of chemical matter under ‘plausible’ prebiotic conditions giving rise to non-enzymatic production of life’s building blocks and their proto-biopolymers, which then somehow morphed into living cells - a process that is usually depicted with question marks (Jortner, 2006). However, all experimental evidence from prebiotic organic syntheses and astrochemical observations show only complex mixtures of carbon-based compounds, including ‘tars’ (Shapiro, 1986), as required by the second law of thermodynamics, with various amounts of biochemical building blocks. The paradigm of complexification of matter - from the simple to the complex - therefore discounts the second law of thermodynamics (the natural diffusional drift in multicomponent solutions and dispersions to ‘homogenize’ mixtures toward greater disorder) and lacks the basic physicochemical mechanisms of how living states could phase-separate (emerge) and keep evolving from inanimate (random) high entropy molecular environments. Hence, there has been a ‘stalling’ progress in origins research because the current fundamental premise of complexification of matter leads to isolated experiments - to Harold’s ‘heap of data’, which lack theoretical framework that would connect them to actual living cells.

Our constraining principles for life’s emergence suggest new experiments with complex chemical mixtures that can represent both prebiotic molecular mixtures, and ‘biotic’ complex mixtures obtained by taking bacterial cells apart. Some potential experiments are briefly described below.

#### Prebiotic complex molecular mixtures

The evolution of historical chemical complexity, viewing early Earth as a ‘giant PCR machine’, can be investigated by building large-scale chemical engineering simulators of prebiotic Earth (Spitzer, 2013). Out of necessity, this is an empirical experimental approach because colloidal phase-separations from multicomponent compositions under cyclic gradients are not theoretically predictable. Nevertheless, enough knowledge has now accumulated to design well-informed empirical experiments based on advances in nano-technology, biotechnology, bacteriology and planetary sciences. The physicochemical model of life’s emergence suggests also limited-scope experiments to address some more specific questions. For instance, biopolymer and metabolite homo-chirality and the cytoplasmic excess of potassium ions are likely physicochemically linked (Spitzer, 2013). This linkage is based on the fact that potassium salts of amino acids crystalize first because of their lower solubility compared to sodium salts; and some of them crystallize as conglomerates - mixed macroscopic crystals of pure enantiomers, e.g., glutamate crystalizes in pure enantiomeric conglomerates (Ault, 2004). The thermodynamics and kinetics of such cyclic precipitations and dissolutions - in multicomponent mixtures in confined microspaces - have not been investigated but they are likely to lead to local amplifications of enantiomeric excess (Weissbuch & Lahav, 2011). A different physicochemical experiment could evaluate which of the chemically evolving building blocks could form coordination complexes with multivalent ions such as Zn^2+(^aq), Fe^2+^(aq), Fe^3+^(aq) and other ions in order to keep them from precipitation in neutral and alkaline pH regions. For instance HCN is widely distributed in the Universe, and the cyanide ion readily forms coordination complexes with ferrous and ferric ions in water, possibly providing a prebiotic redox couple in confined local solutions enriched with proto-metabolite enantiomers and potassium ions. Yet other physicochemical experiments could characterize surface and interfacial properties of ‘intractable tars’ (obtained in Stanley Miller type experiments and during organic non-enzymatic syntheses, and observed as tholins on Saturn’s moon Titan), for their propensity to self-aggregate in hot electrolytes into proto-vesicles or other confining microspaces or matrices.

#### Complex contemporary ‘biotic’ molecular mixtures

The physicochemical basis of meso-evolution and macro-evolution defined earlier involve breakage and fusions of membranes allowing for potential recombinations of segments of DNA from different bacterial species or strains. This mechanism suggests that cyclic manipulations of temperature and dehydration could be developed into a novel method of ‘genetic engineering’ (natural, environmental and evolutionary) without involving divalent cationic salts or electroporation in order to force an (engineered) plasmid through anionic cell envelopes; rather, mixed bacterial populations could be subjected to temperature and dehydration cycles to force membrane fusions concurrently with melting and hybridization of different nucleic acids and thus observe bacterial cellular emergence and evolution in the laboratory. New intermediate cellular structures (the ‘missing links’?) could appear between the surviving and evolving prokaryotic and eukaryotic cellular designs, with a potential to further evolve into a more stable state, pending the properties of the nutrient environment. Presumably, mixed bacterial species that are phylogenetically close may yield ‘new’ bacterial species or bacterial strains, and those that are phylogenetically far apart may merely die off in the process, i.e. become extinct.

While the historical emergence of life was posed by Darwin as the question of the origin of biological species, the contemporaneous emergence of life (spontaneous generation) was disposed of by Pasteur’s experiments with swan neck flasks, resulting in the mantra - ‘a life only from life’ - even at the micron-size microbial level. This is still the standard biological law but it is time to re-visit the question anew (Spitzer, 2014). It is a question of the nature of life, the question of ‘being alive vs. being dead’ - the fundamental pre-condition for Darwinian evolution - a problem that biochemists have been shy to tackle. How could cycling gradients (temperatures, water activities etc.) restructure dead bacterial molecules (in various degrees of separation) into living cells? Or has the extant prokaryotic life evolved to such a very high degree of physicochemical sophistication (over the last 3.5 billion years) that cycling chemical processes are no longer functional with contemporary biopolymers to bring about living states? The inherent instability and plasticity of bacterial genomes (Darmon & Leach, 2014), the physical chemistry of DNA and RNA melting and hybridizations and the protocols of genetic engineering and polymerase chain reactions suggest that cyclic processes may indeed re-assemble ‘dead’ bacterial biomacromolecules into crowded living states - initially with poorly functional cell envelopes arranged in proto-biofilms, with growth (metabolism) powered largely by cycling gradients, and with poorly developed heredity (DNA replication and cell division) - not yet life as we know it; could such a system evolve cyclically in a suitable nutrient medium into one where cellular heredity could be observed?

### Conclusion

While the nature and emergence of life can be sought in unknown biophysical laws as suggested by Schrödinger a long time ago, our elaboration of Atkins’s physicochemical (thermodynamic) view is more directly fertile. Given that on early Earth there were many different complex molecular mixtures behaving according to the physical and chemical laws as we know them today, then the emergence of life proceeded from the high entropy inanimate chemical complexity of Earth’s environments to the lower entropy (phase-separated) colloidal ‘proto-biofilms’, in which proto-nucleic acids, proto-proteins and cell envelopes were cyclically co-evolving. When cell envelopes chemically evolved and sufficiently stabilized, single cell organisms could appear with low rate of horizontal gene transfer and efficient internal homeostasis controlled by membrane proteins, which brought about observable cellular heredity and Darwinian evolution (biology). The basic physicochemical mechanism is the continuous availability of cycling (proto-PCR) energies, which bring about phase separations on a colloidal scale of about 10 - 10,000 nm (typical sizes of living microbes), in a chemical pattern-forming manner, thereby counteracting the unavoidable drift toward randomness as embodied in the second law of thermodynamics. In contrast (and echoing Harold’s view), the current paradigm of ‘complexification of matter’ appears to have run its course: it invokes pre-designed actions and energies of external agents to ‘construct life’ in a non-evolutionary manner, and it does not suggest any physicochemical mechanisms of defeating the diffusional drift to disorder given by the second law of thermodynamics - the de-mixing of non-equilibrium primordial ‘molecular soups’ by the action of non-covalent molecular forces into cellular living states and nutrient environments. Thus Dobzansky’s dictum about the explanatory power of Darwinian evolution for biology can be restated for origins research as: ‘nothing in the origin of life makes sense except in the light of diurnal gradients acting on complex chemical mixtures.

## Acknowledgements

This article is dedicated to late Prof. Robert Shapiro.

We thank Robert Shapiro, Michael Russell, David Deamer, Franklin Harold, Elio Schaechter, George Fox and Gary Pielak for comments on early drafts of these concepts. B.P. work is supported by the NWO TOP-GO (L.10.060), NWO TOP-Punt (718.014.001) grants and an ERC Advanced Grant. J.S. thanks Karel Spitzer (Entomology Institute of Czech Academy of Sciences) for discussions about Darwinian evolution, and MCP Inc. for support.

## References

1. Andersen, O.S. (2015) Perspectives on the response to osmotic challenges. J Gen Physiol 145: 371–372.

2. Atkins, P. (2011) On Being. Oxford: Oxford University Press, pp. 29–30, 102.

3. Ault, A. (2004) The monosodium glutamate story: the commercial production of MSG and other amino acids. J Chem Educ 81: 347–355.

4. Bauermeister, A., Moeller, R., Reitz, G., Sommer, S., and Rettberg, P. (2011) Effect of relative humidity on *Deinococcus radiodurans’* resistance to prolonged desiccation, heat, ionizing, germicidal, and environmentally relevant UV radiation. Microb Ecol 61: 715–22.

5. Bedau, M.A., and Cleland, C.E. (2010) *The Nature of Life: Classical and Contemporary Perspectives from Philosophy and Science*. Cambridge: Cambridge University Press.

6. Benner, S.A. (2010) Defining life. Astrobiology 10: 1021–1030.

7. Bernal, J.D. (1967) *The Origin of Life*. Cleveland: The World Publishing Co., pp. 199–251.

8. Bernstein, M. (2006) Prebiotic materials from on and off the early Earth. Phil Trans R Soc B 361: 1689–1702.

9. Boersma, A.J., Zuhorn, I.S., and Poolman, B. (2015) A sensor for quantification of macromolecular crowding in living cells. Nature Methods 12: 227–9.

10. Bogosian, G., and Bourneuf, E.V. (2001) A matter of bacterial life and death. EMBO Rep 2: 7–704.

11. Bray, D., Levin, M.D., and Morton-Firth, C.J. (1998) Receptor clustering as a cellular mechanism to control sensitivity. Nature 393: 85–88.

12. Bray, D. (2005) Flexible peptides and cytoplasmic gels. Genome Biology 6: 106.

13. Carrasco N., Schmitz-Afonso, I., Bonnet, J.Y., Quirico, E., Thissen, R., Dutuit O., et al. (2009) Chemical characterization of Titan’s tholins: solubility, morphology and molecular structure revisited. J Phys Chem A 113: 11195–11203.

14. Chyba, C., and Sagan, C. (1992) Endogenous production, exogenous delivery and impact-shock synthesis of organic molecules: an inventory for the origins of life. Nature 355: 125–32.

15. Cleland, C.E., and Chyba, C.F. (2002) Defining ‘life’. Orig Life Evol Biosph 32: 387–93.

16. Darmon, E., and Leach, F. (2014) Bacterial genome instability. Microbiol. Mol. Biol. Revs. 78: 1–39.

17. Davey, H.M. (2011) Life, death and in-between: meanings and methods in microbiology. Appl Environ Microbiol 77: 5571–5576.

18. Deamer, D.W., and Fleischaker, G.R. (1994) *Origins of Life: the Central Concepts*. Boston: Jones & Bartlett Publishers, pp.11–12.

19. Dill, K.A. (1990) Dominant forces in protein folding. Biochemistry 29: 7133–7155.

20. Doolittle, W.F. (1999) Phylogenetic classification and the universal tree. Science 284: 2124–2128.

21. Dworkin, J.P., Deamer, D.W., Sandford, A. S., and Allamandola, L.J. (2001) Self-assembling amphiphilic molecules: Synthesis in simulated interstellar/precometary ices. Proc Natl Acad Sci USA 98: 815–819.

22. Ehrenfreund, P., and Cami, J. (2010) Cosmic carbon chemistry: from the interstellar medium to the early Earth. Cold Spring Harb Perspect Biol. http://cshperspectives.cshlp.org/content/2/3/a002097.

23. Ehrenfreund, P., Rasmussen, S., Cleaves, J., and Chen, L. (2006) Experimentally tracing the key steps in the origin of life: the aromatic world. Astrobiology 6: 490–520.

24. Eisenberg, D. (2003) The discovery of α-helix and β-sheet, the principal structural features of proteins. Proc Natl Acad Sci USA 100: 11207–11210.

25. Eisenberg, D., Marcotte, E.M., Xenarios, I., and Yeates, T.O. (2000) Protein function in the post-genomic era. Nature 405: 823–6.

26. Ellis, R.J. (2001) Macromolecular crowding—obvious but underappreciated. Trends Biochem Sci 26: 597–604.

27. England, J.L. (2013) Statistical physics of self-replication. J. Chem. Phys. 139, 121923; doi: 10.1063/1.4818538.

28. Foffi, G., Pastore, A., Piazza, F., and Temussi, P.A. (2013) Macromolecular crowding: chemistry and physics meet biology (Ascona, Switzerland, 10-40 June, 2012). Phys. Biol. 10(4):040301.

29. Frenkiel-Krispin, D., Ben-Avraham, I., Englander, J., Shimoni, E., Wolf, S.G., and Minsky, A. (2004) Nucleoid restructuring in stationary-state bacteria. Mol Microbiol 51: 395–405.

30. Fry, I. (1999) *The Emergence of Life on Earth*. A Historical and Scientific Overview. New Brunswick: Rutgers University Press.

31. Gierasch, L.M., and Gershenson, A. (2009) Post-reductionist protein science, or putting Humpty Dumpty back together again. Nature Chem Biol 5: 774–777.

32. Harold, F.M. (2014) *In Search of Cell History*. Chicago: Chicago University Press, pp. 164, 189.

33. Harold, F.M. (2005) Molecules into cells: specifying spatial architecture. Microbiol Mol Biol Rev 69: 5445–64.

34. Henderson, L.J. (1913) *The Fitness of the Environment*. New York: MacMillan.

35. Hud, N.V., Cafferty, B.J., Krishnamurthy, R. and Williams, L.D. (2013) The origin of RNA and “my grandfather’s axe”. Chem Biol. 20: 466–74.

36. Jortner, J. (2006) Conditions for the emergence of life on the early Earth: summary and reflections. Phil Trans R Soc B 361: 1877–1891.

37. Joyce, G. (1994) Foreword. In *Origins of Life: The Central Concepts*. Deamer, D.W., and Fleischaker, G.R. (eds). Boston: Jones and Bartlett, pp. xi–xii.

38. Keating, C.D. (2012) Aqueous phase separation as a possible route to compartmentalization of biological molecules, Accounts of Chemical Research 45: 2114–2124.

39. Kell, D.B., and Oliver, S.G. (2003) Here is the evidence, now what is the hypothesis? The complementary roles of inductive and hypothesis-driven science in the post-genomic era. BioEssays 26: 99–105.

40. Kim, B.-H., and Gadd, G.M. (2008) *Bacterial Physiology and Metabolism*. Cambridge: Cambridge University Press.

41. Konopka, M.C., Sochacki, K.A., Bratton, B., Shkel, I.A., Record, M.T., and Weisshaar, J.C. (2009) Cytoplasmic protein mobility in osmotically stressed *Escherichia coli*. J Bacteriol 191: 231–237.

42. Koonin, E.V. (2009) Summary and survey: Darwinian evolution in the light of genomics. Nucleic Acids Res 37: 1011–1034.

43. Koonin, E.V. (2011) Carl Woese’s vision of cellular evolution and the domains of life. RNA Biology 11: 197–204.

44. Kornberg, A. (2000) Ten commandments: lessons from the enzymology of DNA replication. J. Bacteriol. 182: 3613–3618.

45. Lahav, N. (1999) *Biogenesis-theories of life’s origins*. Oxford: Oxford University Press.

46. Lane, N. (2015). The Vital Question: Energy, Evolution and the Origins of Complex Life. New York: W.W. Norton & Co., pp. 89–121.

47. Laue, T. (2012) Proximity energies: a framework for understanding concentrated solutions. J Mol Recognit 25:165–73.

48. Loeb, J. (1962) *The Mechanistic Conception of Life*. Cambridge: Harvard University Press, pp. 5–34.

49. Luisi, P.G. (1998) About various definitions of life. Orig Life Evol Biosph 28: 613–622.

50. López, D., and Kolter, R. (2010). Functional microdomains in bacterial membranes. Genes & Dev 24: 1893–1902.

51. Marmur, J., and Doty, P. (1962). Determination of the base composition of deoxyribonucleic acid from its thermal denaturation temperature. J Mol Biol 5: 109–118.

52. Martin, W., Baross, J., Kelley, D., and Russell, M.J. (2008) Hydrothermal vents and the origin of life. Nat Rev Microbiol 6: 805–814.

53. McConkey, E.H. (1982) Molecular evolution, intracellular organization, and the quinary structure of proteins. Proc Natl Acad Sci USA 79: 3236–3240.

54. McDonald, G.D., Khare, B.N., Thompson, W.R., and Sagan, C. (1991) CH_4_/NH_3_/H_2_O spark tholin: chemical analysis and interaction with Jovian aqueous clouds. Icarus 94: 354–67.

55. Mika, J.T., and Poolman, B. (2011) Macromolecule diffusion and confinement in prokaryotic cells. Curr Opin Biotechnol 22: 117–126.

56. Miller, S.L. (1953) A production of amino acids under possible primitive Earth conditions. Science 117: 528–529.

57. Mitchell, P. (1979) Compartmentation and communication in living systems. Ligand conduction: a general catalytic principle in chemical, osmotic, and chemiosmotic reaction systems. Eur J Biochem 95: 1–20.

58. Monteith, W.B., Cohen, R.D., Smith, A.E., Guzman-Cisnerosa, E., and Pielak, G.J. (2015) Quinary structure modulates protein stability in cells. Proc Natl Acad Sci USA. 112: 1739–42.

59. Oliver, J.D. (2010) Recent findings on the viable but non-culturable state in pathogenic bacteria. FEMS Microbiol Rev 34: 415–25.

60. O’Malley, M.A., and Koonin, E.V. (2011) How stands the Tree of Life a century and a half after *The Origin*? Biology Direct 6: 32.

61. Orgel, L.E. (2004) Prebiotic chemistry and the origin of the RNA world. Crit Rev Biochem Mol Biol 39: 99–123.

62. Pal, S. K., Peon, J., Bagchi, B., and Zewail, A. H. (2002) Biological water: femtosecond dynamics of macromolecular hydration. J Phys Chem B 106: 12376–12395.

63. Park, S., Moilanen, D. E., and Fayer, M. D. (2008) Water dynamics—the effects of ions and nanoconfinement. J Phys ChemB 112: 5279–5290.

64. Partikian, A., Ölveczky, B., Swaminathan, R., Li, Y. X., and Verkman, A. S. (1998) Rapid diffusion of green fluorescent protein in the mitochondrial matrix. J Cell Biol 140: 821–829.

65. Pascal, R., and Pross, A. (2015) Stability and its manifestation in the chemical and biological worlds. Chem. Commun., 51: 16160 - 6165.

66. Patel, B.H., Percivalle, C., Ritson, D.J., Duffy, C.D., and Sutherland, J.D. (2015) Common origins of RNA, protein and lipid precursors in a cyanosulfidic protometabolism. Nature Chemistry 7: 301–307.

67. Pauling, L., and Corey, R.B. (1956) Specific hydrogen bond formation between pyrimidines and purines in deoxyribonucleic acids. Archives of Biochemistry and Biophysics 65: 164–181.

68. Parry, B.R., Surovtsev, I.V., Cabeen, M.T., O’Hern, C.S., Dufresne, E.R., and Jacobs-Wagner, C. (2014) The bacterial cytoplasm has glass-like properties and is fluidized by metabolic activity. Cell 156: 183–194.

69. Perutz, M. (1991). Physics and the riddle of life. In Is Science Necessary? Essays on Science & Scientists. Oxford: Oxford University Press, pp. 242–259.

70. Petrov, A.S., Gulen, B., Norris, A.M., Kovacs, N.A., Bernier, C.R., Lanier, K.A., Fox, G.E., Harvey, S.C., Wartell, R.M., Hud, N.V., and Williams, L.D. (2015) “History of the ribosome and the origin of translation”, Proc. Natl. Acad. Sci. U.S.A. 112: 15396–15401.

71. Pizzarello, S., and Shock, E. (2010) The organic composition of carbonaceous meteorites: the evolutionary story ahead of biochemistry. Cold Spring Harb Perspect Biol http://cshperspectives.cshlp.org/content/2/3/a002105.

72. Poolman, B., Spitzer, J.J., and Wood, J.M. (2004) Bacterial osmosensing: roles of membrane structure and electrostatics in lipid-protein and protein-protein interactions. Biochim Biophys Acta 1666: 88–104.

73. Potrykus, K., and Cashel, M. (2008). (p)ppGpp: still magical? Annu Rev Microbiol 62: 35–51.

74. Raulin, F., Brassé, C., Poch, O., and Coll, P. (2012) Prebiotic-like chemistry on Titan. Chem Soc Rev 41: 5380–93.

75. Record, M. T., Courtenay, E. S., Cayley, S., and Guttman, J. H.. (1998) Biophysical compensation mechanisms buffering *E. coli* protein-nucleic acid interactions against changing environments. Trends Biochem Sci 23: 190–194.

76. Rhee, Y.M., Lee, T.J., Gudipati, M.S., Allamandola, L.J., and Head-Gordon, M. (2007) Charged polycyclic aromatic hydrocarbon clusters and the galactic extended red emission. Proc Natl Acad. Sci USA 104: 5274–5278.

77. Rose, G.D., and Wolfenden, R. (1993) Hydrogen bonding, hydrophopbicity, packing and protein folding. Annu Rev Biophys Biomol Struct 22: 381–415.

78. Rothschild, L.J. (2003) The Sun: the impetus of life. In Evolution on Planet Earth: The Impact of the Physical Environment. Rothschild, L. and Lister, A. (eds). London: Academic Press, pp. 87–107.

79. Rowe, A.J. (2011) Ultra-weak reversible protein-protein interactions. Methods 54: 157–166.

80. Saladino, R., Botta, G., Pina, S., Costanzo, C., and Di Mauro, E. (2012) Genetics first or metabolism first? The formamide clue. Chem Soc Rev 41: 5526–5565.

81. Sarkar, M., Smith, A.E., and Pielak, G.J. (2013) Impact of reconstituted cytosol on protein stability. Proc Natl Acad Sci. USA. 110: 19342–19347.

82. Schaechter, M., Ingraham, J.L., and Niedhart, F.C. (2006) Microbe. Washington D.C.: ASM Press.

83. Scharf, C., Virgo, N., Cleaves II, H.J., Aono, M., Aubert-Kato, N., Aydinoglu, A., Barahona, A., et al. (2015) A strategy for origins of life research. Astrobiology 15: 1032–1038.

84. Schildkraut, C., and Lifson, S. (1965). Dependence of the melting temperature of DNA on salt concentration. Biopolymers 3: 195–208.

85. Schrödinger, E. (2012) *What is Life?* Cambridge: Cambridge University Press.

86. Schuster, T.M., and Laue, T.M. (1994) *Modern Analytical Ultracentrifugation: Acquisition and Interpretation of Data for Biological and Synthetic Polymer Systems*. Boston: Birkhäuser.

87. Shapiro, R. (1986) *Origins: a Skeptic’s Guide to the Creation of Life on Earth*. New York: Simon & Schuster.

88. Segré, D., Ben-Eli, D., Deamer, D.W., and Lancet, D. (2001) The lipid world. Orig Life Evol Biosph 31: 119–45.

89. Siegele, D.A., and Kolter, R. (1992) Life after log. J Bacteriol 174: 345–348.

90. Southall, N.T., Dill, K.A., and Haymet, A.D.J. (2002) A view of the hydrophobic effect. J Phys Chem B 106: 521–533.

91. Spitzer, J., Pielak, G., and Poolman, B. (2015) Emergence of life: physical chemistry changes the paradigm. Biology Direct 10: 33. doi:10.1186/s13062-015-0060-y.

92. Spitzer, J. (2014) The continuity of bacterial and physicochemical evolution: theory and experiments. Res Microbiol 165: 457–461.

93. Spitzer, J. (2013) Emergence of life from multicomponent mixtures of chemicals: the case for experiments with cycling physicochemical gradients. Astrobiology 13: 404–413.

94. Spitzer, J., and Poolman, B. (2013) How crowded is the prokaryotic cytoplasm? FEBS Letts 587: 2094–2098.

95. Spitzer, J. (2011) From water and ions to crowded biomacromolecules: in vivo structuring of a prokaryotic cell. Microbiol Mol Biol Revs 75: 491–506.

96. Spitzer, J., and Poolman, B. (2009). The role of biomacromolecular crowding, ionic strength and physicochemical gradients in the complexities of life’s emergence. Microbiol Mol Biol Revs 73: 371–388.

97. Spitzer, J., and Poolman, B. (2005) Electrochemical structure of the crowded cytoplasm. Trends Biochem Sci 30: 536–541.

98. Spitzer, J.J. (2003) Maxwellian double layer forces:? from infinity to contact. Langmuir 19: 7099–7111.

99. Spitzer, J.J. (1984) A re-interpretation of hydration forces. Nature 310: 396–397.

100. Srere, P.A. (1985) The metabolon. Trends Biochem Sci 10: 109–110.

101. Stewart, E.J. (2012) Growing unculturable bacteria. J Bacteriol 194: 4151–60.

102. Stoker, C.R., Boston, P.J., Mancinelli, R.L., Segal, W., Khare, B.N., and Sagan, C. (1990) Microbial metabolism of tholin. Icarus 85:241–56.

103. Stüeken, E.E., Anderson, R.E., Bowman, J.S., Brazelton, W.J, Colangelos-Lillis, J., Goldman, A.D., Som, S.M., and Baross, J.A. (2013) Did life originate from a global chemical reactor? Geobiology http://onlinelibrary.wiley.com/doi/10.1111/gbi.12025/full.

104. Sykes, M.T., and Williamson, J.R. (2009) A complex assembly landscape for the 30S ribosomal subunit. Annu Rev Biophys 38: 197–215.

105. Szostak, J.W., and Deamer, D.W. (eds). (2011) *The Origins of Life*. Cold Spring Harbor: Cold Spring Harbor Laboratory Press.

106. Szostak, J.W. (2012) Attempts to define life do not help to understand the origin of life. J Biomol Struct Dyn 29: 599–600.

107. Tang, T.-Y., Hak, C.R.C., Thompson, A.J., Kuimova, M.K., Williams, D.S., Perriman, A.W. and Mann, S. (2014) Fatty acid membrane assembly on coacervate microdroplets as a step towards a hybrid protocell model. Nature Chemistry 6: 527–533.

108. Theillet, F.-X., Binofli, A., Frembgen-Kesner, T., Hingorani, K., Sarkar, M., Kyne, C., Li, C., Crowley, P.B., Gierasch, L., Pielak, G.J., Elcock, A.H., Gershenson, A., and Selenko, P. (2014) Physicochemical properties of cells and their effects on intrinsically disordered proteins (IDPs). Chem Rev 114: 6661–6714.

109. Traub, P., and Nomura, M. (1968). Structure and function of *E. coli* ribosomes. V. Reconstitution of functionally active 30S ribosomal particles from RNA and proteins. Proc Natl Acad Sci USA. 59: 777–84.

110. Trifonov, E.N. (2012). Definition of life: navigation through uncertainties. J Biomol Struct Dyn 29: 647–650.

111. van den Bogaart, G., Hermans, N., Krasnikov, V., and Poolman, B. (2007) Protein mobility and diffusive barriers in *Escherichia coli*: consequences of osmotic stress. Mol Microbiol 64: 858–71.

112. Wang, Q., Zhuravleva, A., and Gierasch, L.M. (2011) Exploring weak, transient protein-protein interactions in crowded *in vivo* environments by in-cell nuclear magnetic resonance spectroscopy. Biochemistry 50: 9225–9236.

113. Weissbuch, I., and Lahav, M. (2011) Crystalline architectures as templates of relevance to the origins of homochirality. Chem Rev 111: 3236–3267.

114. Woese, C.R. (1998). The universal ancestor. Proc Natl Acad Sci USA 95: 6854–6859.

115. Woese, C.R. (2000). Interpreting the universal phylogenetic tree. Proc Natl Acad Sci USA 97: 8392–8396.

116. Woese, C.R. (2002) On the evolution of cells. Proc Natl Acad Sci USA 99: 8742–8747.

117. Woese, C. R. (2004) A new biology for a new century. Microbiol Mol Biol Rev 68: 173–186.

118. Wong, J. T.-F. (2005) Coevolution theory of the genetic code at age thirty. BioEssays 27: 416–425.

119. Wood, J.M. (2015) Bacterial responses to osmotic challenges. J Gen Physiol 145: 381–388.

120. Zaikowski, L., Friedrich, J.M., and Eldredge, N. (eds). (2008) Chemical Evolution across Space and Time. Washington D.C.: American Chemical Society, ACS Symposium Series 981.

121. Zhou, H.X., Rivas, G., and Minton, A.P. (2008). Macromolecular crowding and confinement: biochemical, biophysical, and potential physiological consequences. Annu Rev Biophys 37: 375–97.

122. Zimmerman, S.B., and Minton, A.P. (1993). Macromolecular crowding: biochemical, biophysical, and physiological consequences. Annu Rev Biophys Biomol Struct 22: 27–65.

